# Psychiatric Genomics Research During the COVID-19 Pandemic: A Survey of Psychiatric Genomics Consortium Researchers

**DOI:** 10.1101/2020.10.08.331421

**Authors:** Jerry Guintivano, Danielle Dick, Cynthia M Bulik

**Affiliations:** Department of Psychiatry, University of North Carolina at Chapel Hill, Chapel Hill, North Carolina, USA; Departments of Psychology and Human & Molecular Genetics, Virginia Commonwealth University, Richmond, Virginia, USA; Department of Medical Epidemiology and Biostatistics, Karolinska Institutet, Stockholm, Sweden; Department of Nutrition, University of North Carolina at Chapel Hill, Chapel Hill, North Carolina, USA

**Keywords:** COVID-19, academic medicine, faculty, promotion, funding

## Abstract

Between April 20, 2020 and June 19, 2020 we conducted a survey of the membership of the Psychiatric Genomics Consortium (PGC) to explore the impact of COVID-19 on their research and academic careers. A total of 123 individuals responded representing academic ranks from trainee to full professor, tenured and fixed-term appointments, and all genders. The survey included both quantitative and free text responses. Results revealed considerable concern about the impact of COVID-19 on research with the greatest concern reported by individuals in non-permanent positions and female researchers. Concerns about the availability of funding and the impact of the pandemic on career progression were commonly reported by early career researchers. We provide recommendations for institutions, organizations such as the PGC, as well as individual senior investigators to ensure that the futures of early career investigators, especially those underrepresented in academic medicine such as women and underrepresented minorities, are not disproportionately disadvantaged by the COVID-19 pandemic.

## Introduction

Since the initial outbreak in December 2019, the novel coronavirus SARS-CoV-2 (COVID-19) has mushroomed into a global pandemic affecting every aspect of life. In an effort to reduce transmission, many governments, universities, and other research institutions issued work from home orders for non-essential workers. Interruptions were widespread on both work and home fronts. Researchers or their family members were infected by COVID-19, schools were closed leaving little time or space for work, and the unpredictability of the course of the pandemic led to persistent anxiety and distress for most people in the world.

Specifically, many clinician-researchers were seconded to COVID-related clinical duties; patient facing research was halted or postponed; many basic science researchers were mandated to halt all laboratory-based activities; and academic medical centers faced enormous financial consequences (Colenda, Applegate, Reifler, & Blazer, 2020; Kim et al., 2020; Weissman, Klump, & Rose, 2020). In person teaching was suspended requiring rapid adaptation to remote teaching platforms. In addition, other academic activities, such as conferences and face-to-face meetings, were cancelled or transitioned to virtual formats, interrupting training, networking, and other means of scientific information exchange. Several studies have documented challenges that have been faced by academics at different career levels, different genders, and different family structures, and concerns have been raised, especially for early-career researchers and women regarding the long-term impact of the pandemic on their career progression (Denfeld et al., 2020).

As a service especially to our early career researchers, the Psychiatric Genomics Consortium (PGC) wanted to better understand the effects of COVID-19 on its members and their research. We modeled our questionnaire after Weissman et al (Weissman et al., 2020) and assessed: (a) the impact of COVID-19 on PGC research now and in the foreseeable future; (b) the level of concern about COVID-19-related disruptions of research and the potential impact of these disruptions on individuals’ careers; (c) strategies that respondents thought to be effective in coping with COVID-19-related research disruptions; and (d) respondents’ suggestions for how the PGC or the field should respond to help researchers move through and beyond the current crisis.

To address these aims, we administered an anonymous survey containing both quantitative and free text responses to PGC researchers. Quantitative questions focused on the perceived impact of COVID-19-related research disruptions. We hypothesized that respondents in secure employment positions (e.g., with tenure or permanent contracts) would report less stress and less concern about potential adverse impact of COVID-19 on their research and career, than respondents at earlier stages in their career (e.g., tenure-track or fixed-term contracts, post-doctoral fellows, graduate students). The primary goal of the qualitative, free text questions was to describe participants’ strategies or suggestions, and highlight issues that they felt were important that we had not addressed. We had no hypotheses regarding the free text responses.

## Methods

### Sample and procedures

All members of the Psychiatric Genomics Consortium (PGC) identified by the consortium listserv were invited to participate in the survey. Invitations were sent through the main PGC listserv, through individual Workgroup listservs, and were promoted during regular PGC videoconferences. The survey was launched on April 20, 2020 and closed on June 19, 2020.

Qualtrics was used to administer and store the survey. This study was approved by the University of North Carolina Institutional Review Board Committee for the Protection of Human Subjects.

### Description of the Survey

This mixed-methods survey was composed of 14 Likert-scale items measuring concern about COVID-19 disruptions (rated from 0 = no concern to 10 = extreme concern) and highest level of stress experienced since the outbreak of the pandemic (0 = no stress to 10 =highest level of stress imaginable). One item asked respondents to characterize the proportion of their PGC-related research that had to be completely shut down due to COVID-19, using six options (from 0 to up to 100% in 20% increments). Three categorical items (yes, no, do not know/does not apply) addressed whether studies had been transferred to online; whether researchers were anticipating making changes to their research practices; and whether their institution had made policy changes in response to COVID-19. Each categorical item was followed by open-ended questions inviting further expansion of answers. Participants could provide up to three open-ended responses when asked about strategies they found most effective for dealing with COVID-19 in terms of their research, changes the PGC research community should make to support researchers during and post-COVID-19, and changes the PGC should make to support researchers during and post-COVID-19. A final open-ended question invited commentary on questions that we did not ask.

Demographic data were collected in a final set of questions. Participants were asked to report their gender (Female, Male, Gender variant/nonconforming, choose not to answer), current position (“Faculty appointment (>5 years post training)”, “Faculty appointment (up to 5 years post training) (Early Career Researcher)”, “Graduate student”, “Post-doctoral fellow”, “Resident”, “Other”), type of position (“A position that could lead to tenure or a permanent contract, but I have not yet reached this status”, “A position that is not in the tenure track or permanent contract system”, “A tenured or permanent contract position”, “other”), department/institution/organization of their primary appointment (Genetics, Government Organization, Medicine [other than psychiatry], Non-academic Hospital or Clinic, Other, Psychiatry, Psychology, Public Health, Research Institute), and the country where they hold their primary research appointment (a drop-down menu of countries in the world).

### Data Analysis

Descriptive statistics were calculated for quantitative items using R (R Core Team, 2020). Chi-square tests and analyses of variance (ANOVAs) were used to test hypotheses that individuals would differ on the basis of type of academic position and gender. For analyses on gender, only male and female respondents were used due to small sample sizes of other respondents. A stringent significance threshold of *p* < 0.005 was used to account for multiple testing throughout the study. In addition, Cohen’s d were used to estimate effect sizes.

Responses to open-ended questions were grouped into thematic categories, as follows. For each open-ended question, the first author independently developed a set of themes to capture the responses. Each theme was required to capture at least three responses to the open-ended question. Because the goal of reporting the open-ended responses was strictly descriptive, we made no attempt at establishing reliability of the coding of major themes.

## Results

### Sample Description

A total of 123 individuals completed the survey, with a majority of respondents being female (n = 66, 53.7%) compared to male (n = 48, 39.0%) and gender variant/nonconforming or unreported (n = 8, 6.5%) (**Table 1**). The majority of respondents more held permanent/tenured positions (n = 69, 56.1%) compared to non-permanent faculty (n = 25, 20.3%), trainees (n = 18, 14.6%) or “other” positions (n = 5, 4.1%). Six respondents (4.8%) did not specify their academic position (**Table 1**).

**Table 1.** Characteristics of respondents.

| **Variable** | **Total Sample** |  |
| --- | --- | --- |
|  | **N** | **%** |
| Gender |  |  |
| Female | 66 | 53.7 |
| Male | 48 | 39 |
| Gender variant/nonconforming or not listed | 9 | 7.3 |
|  | 123 |  |
| Position Held |  |  |
| Tenured Faculty | 69 | 56.1 |
| Non-tenured Faculty | 25 | 20.3 |
| Trainee | 18 | 14.6 |
| Other | 5 | 4.1 |
| Not Listed | 6 | 4.9 |
| Proportion of research shutdown |  |  |
| 0% | 37 | 31.6 |
| up to 20% | 40 | 34.2 |
| up to 40% | 17 | 14.5 |
| up to 60% | 11 | 9.4 |
| up to 80% | 7 | 6 |
| up to 100% | 5 | 4.3 |
| Transition PGC studies to online |  |  |
| Yes | 19 | 16.2 |
| No | 43 | 36.8 |
| N/A (e.g., my research is theoretical; my research is secondary data analyses; etc.) | 54 | 46.2 |
| Future changes to research practice |  |  |
| Yes | 35 | 29.9 |
| No | 33 | 28.2 |
| Too soon to tell | 49 | 41.9 |
| Changes in performance evaluations |  |  |
| Yes | 37 | 31.6 |
| No | 45 | 38.5 |
| Not listed | 35 | 29.9 |

Of those who did report their position, a majority held primary research appointments in psychiatry (n = 63, 53.8%) or genetics (n = 26, 22.2%). The remaining respondents indicated various other departments (medicine, psychology, public health), research institutes, or non-academic hospitals. Academic appointments were most commonly in the United States (n = 49, 41.9%), followed by United Kingdom, n = 10; Germany, n = 7; Sweden, n = 7; Australia, n = 4; Denmark, n = 4; Canada, n = 3; Brazil, n < 3; Greece, n < 3; Italy, n < 3; Mexico, n < 3; Netherlands, n < 3; Spain, n < 3; Switzerland, n < 3; Afghanistan, n < 3; Austria, n < 3; Estonia, n < 3; Japan, n < 3; New Zealand, n < 3; Norway, n < 3; Romania, n < 3; South Africa, n < 3. Ten individuals (8.6%) not report the country of their appointment.

A significantly greater proportion of men (n = 37, 77.0% of male sample) reported holding a permanent appointment compared to women (n = 30, 45.5% of female sample) (*χ*^2^(3, 113) = 14.5, *p* = 0.002).

### Quantitative Data

#### COVID-19 Effects on Research Capacity

As shown in **Table 1**, the largest proportion of participants (34.2%) reported that 20% of their research was shut down due to COVID-19. A total of 31.6% of respondents reported no shutdown of their research, many respondents in this category emphasized that their work was mostly on archived data. A total of 4.3% of respondents reported their research to be totally (up to 100%) shut down.

Among those who reported any research disruptions (n = 80, 68.4% of the total sample), some reported being able to transition their studies to online settings (n = 18, 22.5%), whereas others (n = 26, 32.5%) reported not having the need to move their studies online due to the nature of their work being theoretical or secondary data analyses. Further, the majority of respondents who indicated “up to 80%” or “up to 100%” of their research was shut down due to COVID-19 restriction (n = 12) were more likely to also report they were unable to move their research work to an online setting (n = 10).

#### Research and Career-Related Concerns

**Tables 2 and 3** present research related concerns stratified by academic position (**Table 2**) and gender (**Table 3**). On average, the highest levels (mean ≥ 5.0) of research-related concern were related to cancellation of career opportunities, securing future funding, recruitment, and data collection. Intermediate levels of concern (5.0 > mean ≥ 3.0) were disruptions from having to work from home (both technological and domestic concerns), staffing, transferring teaching/supervision to remote/online, budget, and obtaining institutional approvals. Specific problems related to supply procurement (mean = 2.95) and animal research (mean = 1.27) ranked lowest among respondents. There were no significant differences in research-related concerns between appointment groups. However, when stratified by gender, women report significantly greater concerns regarding domestic issues around disruptions from having to work from home (p = 0.001).

**Table 2.**
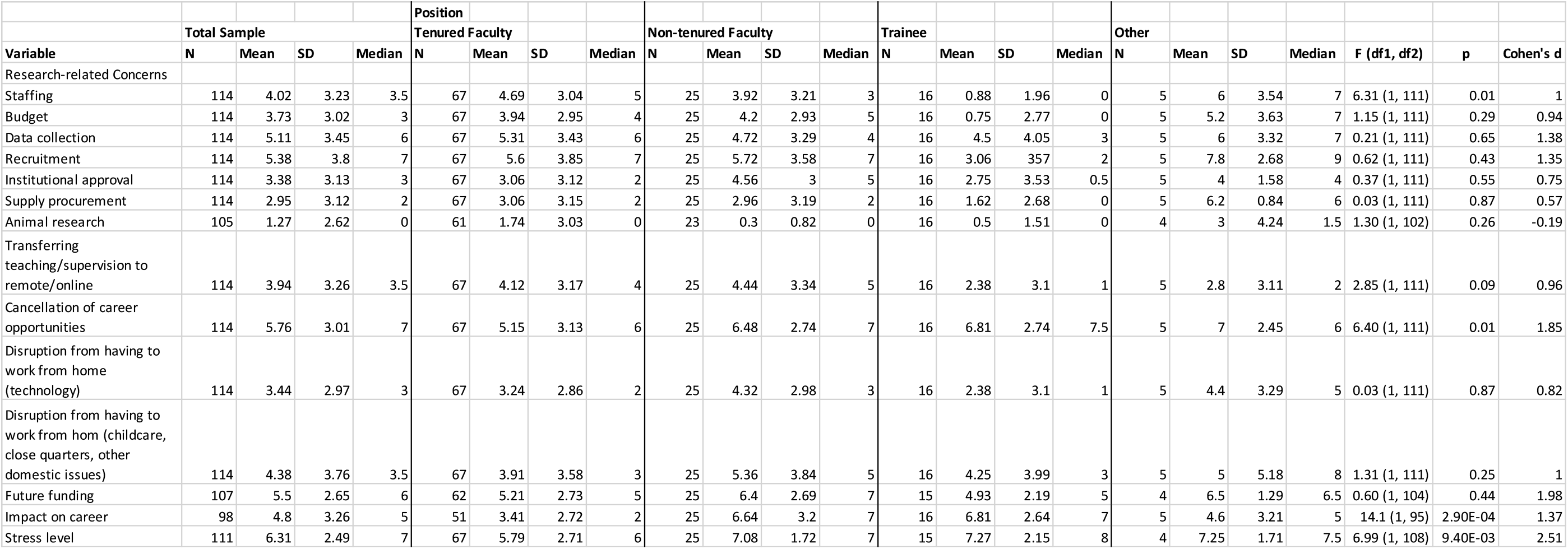
Research-related concerns and stress related to COVID-19 by academic level.

**Table 3.**
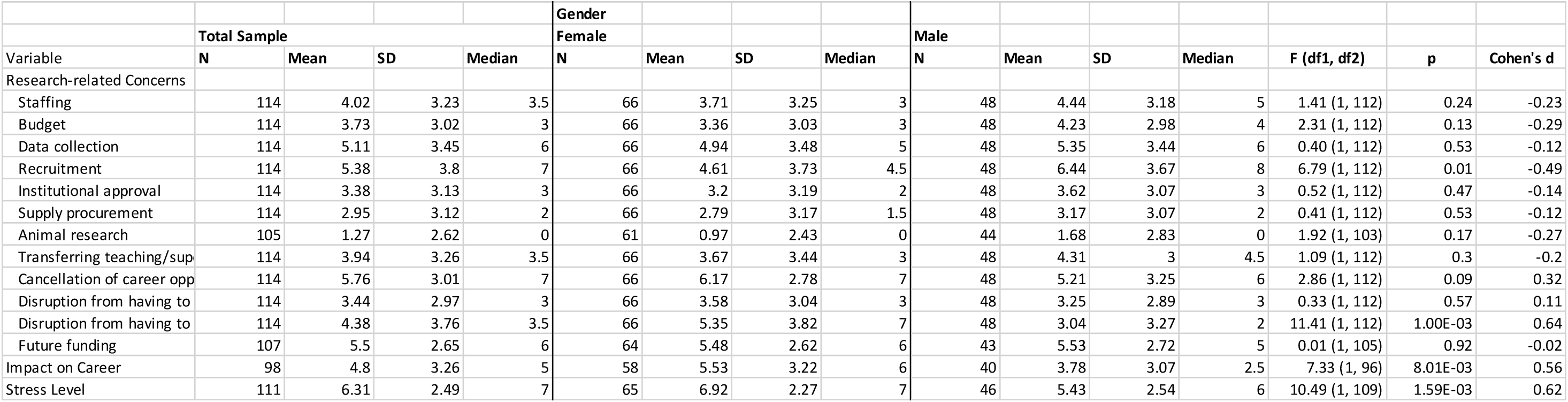
Research-related concerns and stress related to COVID-19 by gender.

Overall, the impact of COVID-19 on career was an intermediate concern (mean = 4.8). However, very large and highly significant differences emerged in career concerns, with trainees and non-tenured faculty reporting higher levels of concern compared to those with permanent positions (p = 2.90E-04, d = 1.37). Moreover, women reported greater levels of career concerns, of medium effect size, compared to men (p = 8.01E-03, d = 0.56).

Stress levels displayed similar patterns as career impact. Stress levels were high among all respondents (mean = 6.3), with higher levels reported among those with non-tenured positions (p = 0.009, d = 2.51) and by females (p = 1.59E-03, d = 0.62).

#### Changes to Future Research Practices

A minority (n = 33, 28.2%) of respondents expected no changes would be made to their research moving forward as a result of COVID-19, while the majority (n = 49, 41.9%) thought it was “too soon to tell” if any changes need to be made (**Table 1**). There was no significant difference across positions regarding the future practices of PGC research (p = 0.007).

#### Institutional Policy Changes

A full 38.5% of respondents reported that there would be no changes in performance evaluations at their institutions related to COVID-19, with approximately one-third of respondents (31.6%) indicated changes, and 29.9% reporting that it was too soon to tell (**Table 1**).

### Qualitative Data

**Table 4** displays findings for the free-text responses, the total number of comments entered for each question, the key themes reflected in those comments, and examples for each of the themes. The examples capture statements in abbreviated and paraphrased form, consistent with our consent statement indicating that we would summarize responses rather than provide specific individual examples.

**Table 4.** Responses to free text questions.

| Table 4. Responses to free text questions |  | Number of comments recorded and themes and illustrative (shortened, paraphrased) Statements identified by open-ended questions. |
| --- | --- | --- |
| *Please share your 1–3 most effective strategies for dealing with COVID-19 in terms of your PGC research. (125 comments)* |  |  |
| 1. Maintain team dynamics (41 / 125, 32.8%) |  |  |
| Utilizing videoconferencing for regular team meetings, being flexible with deadlines, use clear communication |  |  |
| 2. Maintain good personal habits (34 / 125, 27.2%) |  |  |
| Keeping in mind productivity may be reduced, practicing self-care, keeping work and personal areas separate |  |  |
| 3. Reprioritizing research goals (26 / 125, 20.8%) |  |  |
| Spend more effort on dry-lab projects rather than wet-lab, use available time to complete analyses or manuscripts, utilizing existing data for new projects |  |  |
| 4. Shifting recruitment to online means (11 / 125, 8.0%) |  |  |
| Phone interviews rather than face-to-face, development of online recruitment and consent protocols |  |  |
| Please share your 1–3 most effective strategies for transitioning your PGC research to online settings. (Respondents were instructed to skip the question if they had not made such a transition) (24 comments) |  |  |
| 1. Use technology to maintain contact with research participants (10 / 24, 41.7%) |  |  |
| Using online questionnaires and other remote means for recruitment and consent |  |  |
| 2. Use technology to support activities of the research team (8 / 24, 33.3%) |  |  |
| Establishing connections to remote databases, regular teleconferencing with colleagues |  |  |
| Please describe 1–3 changes you expect to make in your PGC research practices as a result of COVID-19. (Respondents were instructed to skip the question if they did not anticipate making changes or had checked “It’s too soon to tell”) (39 comments) |  |  |
| 1. Moving to remote recruitment (10 / 39, 25.6%) |  |  |
| Phone/online recruitment, saliva rather than blood sample collection |  |  |
| 2. Increased use of virtual meetings (9 / 39, 23.1%) |  |  |
| Video conferencing for in lieu of lab or scientific meetings |  |  |
| 3. Organizational changes (6 / 39, 15.4%) |  |  |
| Expectations for decreased budget, reduced personnel |  |  |
| 4. Increased protective measures when collecting samples in-person (5 / 39, 12.8%) |  |  |
| Increased PPE for blood draws |  |  |
| 5. Shifting research priorities (4 / 39, 10.3%) |  |  |
| Add COVID-19 as a research focus |  |  |
| Please describe 1–3 changes the PGC should make to support PGC researchers during and after COVID-19. (74 comments) |  |  |
| 1. Continue to support remote meetings (13 / 74, 17.6%) |  |  |
| Virtual World Congress of Psychiatric Genetics annual meeting, worldwide lab meetings |  |  |
| 2. Facilitate more secondary analyses (12 / 74, 16.2%) |  |  |
| Making data access easier, increasing time frame on existing proposals, prioritize data access for junior researchers |  |  |
| 3. Offer greater support to early career researchers (11 / 74, 14.9%) |  |  |
| Advocate for junior researchers, create support groups for early career researchers |  |  |
| 4. Provide support for researchers directly impacted by COVID-19 (8 / 74, 10.8%) |  |  |
| Leniency for those who see patients and other caregivers, support career advancement for those directly affected by COVID-19 |  |  |
| 5. Provide funding to support researchers (8 / 74, 10.8%) |  |  |
| Bridge funding to sustain PGC research especially among junior researchers and those not part of core PGC funding |  |  |
| 6. Provide online training (6 / 74, 8.1%) |  |  |
| Statistical genetics and other key topic courses |  |  |
| Is there a question about COVID-19’s impact on your PGC research that we should have asked but didn’t? (17 comments) |  |  |
| 1. We should have asked questions regarding clinical researchers and how their practice has shifted to support COVID-19 patients (3 / 17, 17.6%) |  |  |
| 2. Respondents wanted more questions into how broader shutdowns (e.g. school closings, unemployment, work from home) have affected productivity, specifically childcare (3 / 17, 17.6%) |  |  |

#### Effective strategies for dealing with COVID-19-related research challenges

Respondents were asked to provide “1-3 effective strategies for dealing with COVID-19 in terms of your PGC research,” which yielded 125 responses. Four main themes characterized the comments: maintain team dynamics (e.g., utilizing videoconferencing for regular team meetings, being flexible with deadlines, use clear communication) (32.8% of responses); maintain good personal habits (e.g., keeping in mind productivity may be reduced, practicing self-care, keeping work and personal areas separate) (27.2%); reprioritize research goals (e.g., spending more effort on dry-lab projects rather than wet-lab, using available time to complete analyses or manuscripts, utilizing existing data for new projects) (20.8%); and shift recruitment to online approaches (e.g., phone interviews rather than face-to-face, development of online recruitment and consent protocols) (8.0%).

#### Effective strategies for transitioning to online settings

Two themes emerged from the 24 responses to the prompt, “effective strategies for transitioning your PGC research to online settings.” Both themes employ technological approaches to: maintain contact with research participants (e.g., using online questionnaires and other remote means for recruitment and consent) (41.7%) and support activities of the research team (e.g., establishing connections to remote databases, regular teleconferencing with colleagues) (33.3%).

#### Anticipated changes to PGC research practices post-COVID-19

The 39 responses to the prompt “describe 1-3 changes you expect to make in your PGC research practices” could be grouped into five themes: move to remote recruitment (e.g., phone/online recruitment, saliva rather than blood sample collection) (25.6%); increased use of virtual meetings (e.g., video conferencing for in lieu of lab or scientific meetings) (23.1%); organizational changes (e.g., expectations for decreased budget, reduced personnel) (15.4%); increased protective measures when collecting samples in-person (e.g., increased PPE for blood draws) (12.8%); and shifting research priorities (e.g. add COVID-19 as a research focus) (10.3%).

#### Changes PGC should make to support research during and after COVID-19

When asked to “describe 1-3 changes the PGC should make to support PGC researchers during and after COVID-19” 74 responses were provided. These comments fell into six common themes: continue to support remote meetings (e.g. virtual World Congress of Psychiatric Genetics annual meeting, worldwide lab meetings) (17.6%), facilitate more secondary analyses (e.g. making data access easier, increasing time frame on existing proposals, prioritize data access for junior researchers) (16.2%), offer greater support to early career researchers (e.g. advocate for junior researchers, create support groups for early career researchers) (14.9%), provide support for researchers directly impacted by COVID-19 (e.g. leniency for those who see patients and other caregivers, support career advancement for those directly affected by COVID-19) (10.8%), provide funding to support researchers (e.g. bridge funding to sustain PGC research especially among junior researchers and those not part of core PGC funding) (10.8%), and provide online training (e.g. statistical genetics and other key topic courses) (8.1%).

#### Institutional policy changes regarding performance evaluation of researchers

Of the 29 answers to the prompt, “Briefly describe your institution’s policy changes about performance evaluations of researchers,” by far, the most reported change was an extension of performance evaluation periods (65.5% of responses), specifically extending the tenure clock by one year.

### Questions the survey should have asked

Seventeen comments were made in response to the question “Is there a question about COVID-19’s impact on your PGC research that we should have asked but didn’t?” There were two themes that met a minimum of three comments per theme. First, respondents indicated that we should have asked questions regarding clinical researchers and how their practice has shifted to support COVID-19 patients (17.6% of responses). Second, respondents wanted more questions into how broader shutdowns (e.g., school closings, unemployment, work from home) have affected productivity, specifically childcare (17.6% of responses).

## Discussion

Overall, our survey revealed high stress and concern about the impact of COVID-19 on their careers especially in individuals with non-permanent positions and in women. Our results are consistent with those reported across various academic fields (Andersen, Nielsen, Simone, Lewiss, & Jagsi 2020; Brubaker, 2020; Denfeld et al., 2020; Kibbe, 2020; Weissman et al., 2020), and highlight steps that can and should be taken to ensure the ability of early career researchers and female academics not only to survive but to thrive post-pandemic.

Results of the PGC survey align with other surveys reported in the literature that reveal the disproportionate impact that COVID-19-related interruptions have had on female researchers. From having primary responsibility for childcare at home while trying to work from home to concerns about an advancing tenure clock when their productivity is hampered by pandemic-related disruptions, women do appear to be more stressed and more directly impacted than men. This augments the already disproportionate burden of domestic and emotional labor shouldered by female academics (Brubaker, 2020; Jolly et al., 2014; Rao, 2019). Other studies confirm this observation including a disproportionate number of male first authors in papers submitted to journals on COVID-19 (Andersen et al., 2020), journal submissions and productivity in general (Viglione, 2020), and projections of serious interruptions of career progress for women that could adversely affect progress toward gender equity in academe (Sheikh et al., 2018).

A surprising number of institutions were not intending to make allowances on performance evaluations or tenure clocks. This is of significant concern, especially given the documented differential burden placed on junior female faculty in their childbearing years who are entrusted with the majority of childcare duties (Jolly et al., 2014). Senior mentors, institutions, and scientific organizations like the PGC should actively develop and deploy measures to support the careers of junior researchers and those of all genders who have had to take on additional child- or elder-care burdens during this time.

Likewise, although we did not assess ancestry, it has been widely documented in the United States that individuals from underrepresented minority groups have been disproportionately affected by COVID-19 (Moore et al., 2020), which means that not only might more minority researchers be directly affected by COVID-19, but they are also more likely to have connections in socially vulnerable communities and have family members and members of their communities impacted (Nayak et al., 2020). This can divert both time and emotional energy away from career progress and further perpetuate existing systemic inequities in academe (Davis & Fry, 2019).

In many years to come, evaluations of productivity and hiring decisions should explicitly address and account for disruptions encountered during this time. Although many institutions have implemented a single-year extension of tenure clocks, that may not suffice given the prolonged nature of the pandemic. Applications should include the opportunity to describe the impact of the pandemic on an applicant’s life such that it can be factored into the evaluation of the candidate. It is critically important that COVID-19 not set back progress toward equity in science and academe in general, but definitive action must be taken in order to ensure that outcome.

Some caveats and limitations should be considered when interpreting the results. First, the extent to which our sample represents the larger PGC is unknown as we are unable to calculate response rate or representativeness as the survey link was shared widely across PGC groups and subgroups. The composition of the sample, namely primarily female (53.7% of total sample) and in a permanent/tenured position (56.1%), does not necessarily reflect the overall composition of the PGC and may reflect selective participation. Second, given our goal of providing strictly descriptive results, we did not undertake formal efforts at establishing a coding scheme for the free text responses. Third, the survey was deployed relatively early in the pandemic when it was not yet clear how long the disruption to research would go on. Responses could change as the duration of the home- and work-related disruption continues and researchers become increasingly fatigued by the pervasive and persistent disruption. Finally, our failure to assess race and ethnicity was a missed opportunity to capture specific concerns faced by researchers from underrepresented minority groups. As a field, genetics already has considerable underrepresentation of researchers from diverse ancestral backgrounds; our findings for other historically disadvantaged groups (e.g., women, early career investigators) suggest that the pandemic may further exacerbate this underrepresentation.

Recurring themes that emerged focused on the cancellation of career opportunities in terms of networking, but also the financial impact of COVID-19 on job availability as many institutions have implemented hiring freezes. This along with personal economic instability, and concerns about the availability of sources of future research funding lead many researchers to question their future job prospects and the viability of remaining in academe. Many respondents expressed desire for the PGC and senior investigations to devise ways to help boost productivity and success in publications and grant applications—basically devoting greater energy to ensuring the success of early career researchers during this time. The disproportionate underrepresentation of women at higher academic ranks (Carr et al., 2018) is a known phenomenon in many fields of academic medicine, especially for women. It is a critical juncture to ensure that we can shore up promising young investigators such that we can retain them in science and not erase the albeit slow and incremental advances that we have seen in striving for equity in academe (Wingard, Trejo, Gudea, Goodman, & Reznik, 2019).

## Acknowledgements

Dr. Jerry Guintivano is supported by K01 MH116413

Dr. Danielle Dick is supported by NIH R01 AA015416 (Finnish Twin Study), P50 AA022537 (Alcohol Research Center), R25 AA027402 (VCU GREAT), R34 AA027347 (Personalized Risk Assessment), and U10 AA008401 (COGA) from the National Institute on Alcohol Abuse and Alcoholism (NIAAA), and by R01 DA050721 (Externalizing Consortium) from the National Institute on Drug Abuse (NIDA).

Dr. Cynthia Bulik is supported by R01MH120170, R01MH119084, R01MH118278, R34MH113681, R21MH115397, R01MH105684, U01 MH109528, and H79 SM081924. She also acknowledges funding from the Swedish Research Council (Vetenskapsrådet, award: 538-2013-8864).

## References

Andersen, J., Nielsen, M., Simone, N., Lewiss, R., & Jagsi, R. (2020). COVID-19 medical papers have fewer women first authors than expected. Elife., e58807. doi:doi:10.7554/eLife.58807

Brubaker, L. (2020). Women physicians and the COVID-19 pandemic. Jama, 324(9), 835–836. doi:10.1001/jama.2020.14797

Carr, P. L., Raj, A., Kaplan, S. E., Terrin, N., Breeze, J. L., & Freund, K. M. (2018). Gender differences in academic medicine: Retention, rank, and leadership comparisons from the national faculty survey. Acad Med, 93(11), 1694–1699. doi:10.1097/ACM.0000000000002146

Colenda, C. C., Applegate, W. B., Reifler, B. V., & Blazer, D. G., 2nd. (2020). COVID-19: Financial stress test for academic medical centers. Acad Med, 95(8), 1143–1145. doi:10.1097/ACM.0000000000003418

Davis, L., & Fry, D. (2019). College faculty have become more racially and ethnically diverse, but remain far less so than students. Retrieved from https://pewrsr.ch/2GCDVDZ

Denfeld, Q., Erickson, E., Valent, A., Villasana, L., Zhang, Z., Myatt, L., & Guise, J.-M. (2020). COVID-19: Challenges and Lessons Learned from Early Career Investigators. Journal of Women’s Health, 29(6), 752–754. doi:10.1089/jwh.2020.8552

Jolly, S., Griffith, K. A., DeCastro, R., Stewart, A., Ubel, P., & Jagsi, R. (2014). Gender differences in time spent on parenting and domestic responsibilities by high-achieving young physician-researchers. Ann Intern Med, 160(5), 344–353. doi:10.7326/M13-0974

Kibbe, M. R. (2020). Consequences of the COVID-19 Pandemic on Manuscript Submissions by Women. JAMA Surg. doi:10.1001/jamasurg.2020.3917

Kim, C. S., Lynch, J. B., Cohen, S., Neme, S., Staiger, T. O., Evans, L., … Dellit, T. H. (2020). One Academic Health System’s Early (and Ongoing) Experience Responding to COVID-19: Recommendations From the Initial Epicenter of the Pandemic in the United States. Acad Med, 95(8), 1146–1148. doi:10.1097/ACM.0000000000003410

Moore, J. T., Ricaldi, J. N., Rose, C. E., Fuld, J., Parise, M., Kang, G. J., … Territorial Response, T. (2020). Disparities in incidence of COVID-19 among underrepresented racial/ethnic groups in counties identified as hotspots during June 5-1, 2020 - 22 atates, February-June 2020. MMWR Morb Mortal Wkly Rep, 69(33), 1122–1126. doi:10.15585/mmwr.mm6933e1

Nayak, A., Islam, S. J., Mehta, A., Ko, Y. A., Patel, S. A., Goyal, A., … Quyyumi, A. A. (2020). Impact of social vulnerability on COVID-19 incidence and outcomes in the United States. medRxiv. doi:10.1101/2020.04.10.20060962

Rao, A. (2019). Even Breadwinning Wives Don’t Get Equality at Home. The Atlantic.

Sheikh, M. H., Chaudhary, A. M. D., Khan, A. S., Tahir, M. A., Yahya, H. A., Naveed, S., & Khosa, F. (2018). Influences for gender disparity in academic psychiatry in the United States. Cureus, 10(4), e2514. doi:10.7759/cureus.2514

Viglione, G. (2020). Are women publishing less during the pandemic? Here’s what the data say. Nature, 581(7809), 365–366. doi:10.1038/d41586-020-01294-9

Weissman, R. S., Klump, K. L., & Rose, J. (2020). Conducting eating disorders research in the time of COVID-19: A survey of researchers in the field. Int J Eat Disord, 53(7), 1171–1181. doi:10.1002/eat.23303

Wingard, D., Trejo, J., Gudea, M., Goodman, S., & Reznik, V. (2019). Faculty equity, diversity, culture and climate change in academic medicine: A longitudinal study. J Natl Med Assoc, 111(1), 46–53. doi:10.1016/j.jnma.2018.05.004

